# Differing genetics of saline and cocaine self-administration in the hybrid mouse diversity panel

**DOI:** 10.1101/2024.12.04.626933

**Authors:** Arshad H. Khan, Jared R. Bagley, Nathan LaPierre, Carlos Gonzalez-Figueroa, Tadeo C. Spencer, Mudra Choudhury, Xinshu Xiao, Eleazar Eskin, James D. Jentsch, Desmond J. Smith

**Author notes:** Cedars-Sinai Medical Center, 8700 Beverly Blvd, Los Angeles, CA 90048. Department of Pharmaceutical Sciences, Binghamton University, Binghamton, NY 13902. Department of Human Genetics, University of Chicago, Chicago, IL 60637. Sanford Burnham Prebys, La Jolla, CA 92037. To whom correspondence should be addressed: Department of Molecular and Medical Pharmacology, David Geffen School of Medicine, UCLA Box 951735, 23-151A CHS, Los Angeles, CA 90095-1735. (JDJ) (CG-F), (TCS), (XX).

## Abstract

To identify genes that regulate the response to the potentially addictive drug cocaine, we performed a control experiment using genome-wide association studies (GWASs) and RNA-Seq of a panel of inbred and recombinant inbred mice undergoing intravenous self-administration of saline. A linear mixed model increased statistical power for analysis of the longitudinal behavioral data, which was acquired over 10 days. A total of 145 loci were identified for saline compared to 17 for the corresponding cocaine GWAS. Only one locus overlapped. Transcriptome-wide association studies (TWASs) using RNA-Seq data from the nucleus accumbens and medial frontal cortex identified *5031434O11Rik* and *Zfp60* as significant for saline self-administration. Two other genes, *Myh4* and *Npc1*, were nominated based on proximity to loci for multiple endpoints or a *cis* locus regulating expression. All four genes have previously been implicated in locomotor activity, despite the absence of a strong relationship between saline taking and distance traveled in the open field. Our results indicate a distinct genetic basis for saline and cocaine self-administration, and suggest some common genes for saline self-administration and locomotor activity.

## INTRODUCTION

Cocaine misuse, the taking of cocaine for recreational purposes, can lead to cocaine use disorder (CUD), defined as drug taking that leads to significant impairment or distress [1]. Both behaviors represent a significant socioeconomic burden. In the US, over 2 million individuals use cocaine more frequently than once a month [2–6]. Deaths due to overdose in 2018 were 4.5 per 100,000 standard population [4]. Cocaine is a psychostimulant that acts by inhibiting the dopamine transporter, increasing the amount of dopamine in the synaptic cleft [7]. The resulting activation of the mesolimbic dopaminergic pathway, and in particular the nucleus accumbens, play key roles in the reinforcing effects of cocaine [3, 6]. The prefrontal cortex has an important additional function in impulse control and inhibiting drug-induced reinstatement.

Cocaine misuse in humans has broad and narrow sense heritabilities of ∼0.32–0.79 and ∼0.27-0.30, respectively [2]. However, employing genome-wide association studies (GWASs) to identify genes for the disorder in humans has been difficult [3, 5, 6]. The obstacles include recruitment of properly ascertained subjects, interactions between genes and environment, the complex multifactorial nature of cocaine use disorder, and comorbidities between the condition and other psychiatric illnesses [3, 6].

Fully validated models of cocaine use disorder in animals likely await a more complete understanding of mechanisms of this disease [8]. However, intravenous self-administration (IVSA) is regarded as one of the best models of cocaine use disorder in animals, with both construct and predictive validity [9]. We previously used an array of inbred and recombinant inbred mice called the hybrid mouse diversity panel (HMDP) to identify genes involved in four endpoints of cocaine IVSA over a 10 day testing period [10, 11]. The endpoints were the number of cocaine infusions earned, number of active lever presses regardless of whether an infusion was earned, percentage of active lever presses and number of inactive lever presses.

The HMDP consists of ∼30 inbred and 70 recombinant inbred mouse strains that can be used for association mapping of complex traits, including behaviors such as feeding and learning and memory [10, 12, 13, 11, 14–16]. High resolution genetic mapping is facilitated by the inbred strains which possess many recombinations, while the recombinant inbred strains increase resolution but also add statistical power. The HMDP is genetically stable, permitting ascertainment of multiple biological and molecular phenotypes and providing ever more powerful insights. As a control, and to enable studies of differential gene expression, a parallel experiment to the cocaine investigation was performed using saline, with all other factors (handling, surgery, behavioral testing) being identical.

Four sets of observations supported differing genetic causes of cocaine and saline IVSA in the HMDP [10, 11]. First, though we found evidence that cocaine served as a more effective behavioral reinforcer than saline, individual strains engaged in levels of saline self_-_administration that varied considerably relative to cocaine. Second, the behavioral endpoints were much more highly correlated within than between infusates. Third, both narrow and broad sense heritabilities were significantly higher for saline IVSA (∼0.31 and ∼0.44, respectively) than cocaine (∼0.20 and ∼0.32, respectively). Fourth, although neither infusate showed significant loci for individual days, the corresponding genome scans for cocaine and saline segregated nearly completely when subjected to unsupervised clustering.

To increase the statistical power of the cocaine GWASs, we took advantage of the longitudinal data using a linear mixed model that employed fixed and random effects of testing day as a continuous variable, while correcting for population structure using genetic relatedness. A total of 15 unique significant cocaine loci were identified. To further increase statistical power, we used transcriptome-wide association studies (TWASs) to combine the longitudinal genome scans with RNA-Seq data from the nucleus accumbens (NAc) and medial frontal cortex (mFC) of the cocaine cohort. Both the TWASs and GWASs highlighted a number of genes that may be relevant cocaine use disorder [17].

In this report, we use both longitudinal GWASs and TWASs to identify genes for saline-taking and better understand the genetic basis for the differences in cocaine and saline IVSA.

## 2 METHODS

### 2.1 Saline and cocaine intravenous self-administration

A total of 479 and 477 mice from 84 strains of the HMDP completed cocaine and saline IVSA testing respectively, as described [10, 11]. The evaluated strains consisted of 32 inbred and 52 recombinant inbred strains (Table S1). There were 36 testing cohorts, with each cohort featuring a concurrent 13.3 ± 1.1 mice treated with saline and 13.7 ± 1.0 mice with cocaine. To avoid confounds, cohort was used as a nominal covariate in all behavioral analyses.

Animals were obtained from the Jackson Laboratory (Bar Harbor ME) with emplaced indwelling jugular catheters. Catheters were maintained during acclimation by flushing with 0.05 ml of sterile saline and then 0.01 ml heparin solution (500 units ml^-1^) at least once every 3 days, and daily during testing. Catheter patency was confirmed by infusing 0.02 ml propofol (10 mg ml^-1^) followed by reversible loss of muscle tone. The catheters were tested 3 to 4 days before the start of IVSA and after the final session. The final patency rate was 95.1%. Overall attrition rate due to all causes was 9.9%.

For both saline and cocaine, the goal numbers were 3 males and 3 females for each strain and infusate. The realized number (mean ± standard error of the mean, s.e.m.) of mice per strain was 5.7 ± 0.1 for saline (males, 2.9 ± 0.1; females, 2.8 ± 0.1) and 5.7 ± 0.1 for cocaine (males, 2.9 ± 0.1; females, 2.9 ± 0.1) (Table S1). The age of the mice was 11.3 ± 0.1 weeks and 11.4 ± 0.1 weeks for saline and cocaine, respectively. The numbers used are estimated to provide > 80% power to identify a quantitative trait locus (QTL) with an effect size of 10% in the 100 strains of the HMDP [18].

Mice were individually housed and evaluated in the light phase of a 12 h/12 h cycle. Testing took place over 10 consecutive daily sessions using a fixed-ratio-1 (FR1) schedule of reinforcement. Mouse experiments followed all relevant regulatory standards and were approved by the Binghamton University Institutional Animal Care and Use Committee. Analysis of mice from the same strain was distributed across multiple chambers (55.69 × 38.1 × 35.56 cm, MED-307W-CT-D1, Med Associates, VT). The active infusion lever (right or left side of the box) was selected by counterbalancing across strains and sex.

Testing chambers had two response levers, one of which, when pressed, produced an infusion of cocaine or saline depending on cohort, the other of which was inactive. Testing began with illumination of five stimulus lights on the back wall of the chamber and activation of white noise. Priming infusions were not delivered. To encourage conditioning, infusion of the agent was accompanied by a visual cue (flashing of the house light) and extinction of the aperture lights for 20 s. During this time-out period, active lever presses were recorded but delivered no infusions. Pressing the inactive lever had no programmed effect. Testing was for 2 h or until 65 infusions were given, whichever came first. Sterile saline was 0.84 mg ml^-1^ of body weight per infusion and free base cocaine was 0.5 mg kg^-1^. Volumes were 0.67 ml kg^-1^ infusion^-1^ for both substances.

The endpoints analyzed were number of infusions, number of active lever presses, percentage of active lever presses and number of inactive lever presses. The first three endpoints evaluate the propensity for saline self-administration. Active lever presses are the sum of productive lever presses that lead to infusions as well as futile lever presses during the time-out period. Percent active lever presses control for locomotor activity by normalizing active lever presses to total lever presses (active plus inactive). Inactive lever presses may reflect a number of factors, including locomotor activity and the aversive properties of the infusate.

### 2.2 RNA-Seq

RNA-Seq was performed on NAc (core and shell) and mFC of 41 strains exposed to either cocaine or saline IVSA, consisting of 28 inbred and 13 recombinant inbred strains (Table S1) [10, 11]. Tissues were dissected 24 h after the final IVSA test session and samples homogenized in 1 ml of QIAzol reagent (Qiagen, Germantown, MD). RNA was purified using the RNeasy Mini Kit (Qiagen) and abundance measured using NanoDrop (ThermoFisher, Carlsbad, CA). Four strains (A/J, AKR/J, LP/J and NOD/ShiLtJ) were analyzed using individual mice, consisting of 3 males and 3 females for each strain. The remaining samples were pooled by sex to save library construction costs, and consisted of 3 individuals of each sex per strain.

Libraries were created using 100 ng of RNA and TruSeq Stranded RNA kits (Illumina, San Diego, CA). Sequencing of the 392 samples from 41 HMDP strains used an Illumina HiSeq3200 machine. Ten samples per lane were employed with each barcoded sample distributed over multiple lanes to minimize batch effects.

A total of 72.8 ± 0.8 million (M) reads (mean ± s.e.m.) of 75 bp paired-ends were obtained per region per strain averaged across infusates (NAc saline, 72.2 ± 1.5 M; mFC saline, 73.8 ± 1.5 M; NAc cocaine, 70.7 ± 1.3 M; mFC cocaine, 74.6 ± 1.6 M). Samples had excellent quality, with a median FastQC quality score of 40 [19]. STAR aligner was employed to map reads to mouse genome sequence build GRCm38.p6 downloaded from Ensembl96 and quantitated using `htseq-count`, as described [10, 13, 20]. Transcripts with ≥ 6 reads and transcripts per million (TPM) > 0.1 in ≥ 20% of samples for each infusate (cocaine or saline) and brain region (NAc or mFC) were conditional quantile normalized and used for GWAS [21].

Read mapping of spliceforms used STAR and `htseq-count`. Percentage spliced in (psi, or ψ) was calculated as the proportion of transcripts possessing a particular exon [11, 22]. Exons with non-zero standard deviation of ψ and ≥ 5 reads in all samples for each infusate and brain region were retained. Exons with the highest standard deviation of ψ between individuals calculated at the exon level were chosen for each transcript and quantile normalized.

RNA editing sites were quantitated by aligning reads using `HISAT2` v.2.0.4, with realignment of unmapped reads to evaluate hyper-edited sites, as described [11]. Downstream processing steps used the REDIportal database [23]. Editing sites were retained when ≥ 10% of samples in each infusate and brain region had data. The nucleotide changes caused by RNA editing result in decreased read alignments. To quantify the diminished alignment frequencies, we defined the ascertainment rate as the mean proportion of samples with detectable editing of identified sites. All editing sites were A to I. Quantile normalized editing ratios were used for mapping.

### 2.3 Statistical analyses

The software used included `eCAVIAR`, `FaST-LMM`, `FOCUS`, `FUSION`, `GMMAT`, `heritability`, `HISAT2`, `htseq-count`, `lme4`, `LocusZoom`, `R` and `STAR` [20, 24–34]. Standard errors of the mean (s.e.m.) are used throughout.

#### 2.3.1 Behavioral time courses

Saline IVSA endpoints were normalized using rank-based inverse normal transformation (Blom’s method) of individual days, with tied values replaced by their mean. Significance testing of behavioral endpoint time courses used fixed effects of testing day as a continuous variable, sex, active lever (left vs right), testing box, cohort, age and a random intercept and slope for testing day of individual mice.

#### 2.3.2 Mapping loci for IVSA

QTLs for normalized behavioral endpoints were mapped as described using a linear mixed model in `GMMAT` [11, 28]. The model included fixed and random effects of testing day as a continuous variable for individual animals and corrected for population structure via a genetic relatedness matrix. Covariates were sex, active lever (left or right), testing chamber, cohort and age. The model was:

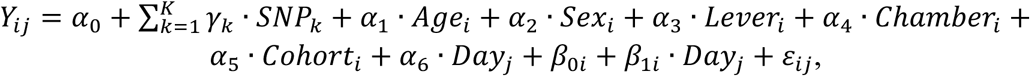

where *Y*_*ij*_ is the normalized saline self-administration endpoint for individual *i* on day *j*, *α*_0_ is the overall intercept, *γ*_*k*_ is the fixed effect for each single nucleotide polymorphism (SNP) *k* (for *k* = 1,…, *K*), *α*_1_, *α*_2_, *α*_3_, *α*_4_, *α*_5_ and *α*_6_ are the fixed effects for age, sex, lever position, chamber, cohort, and day, respectively; 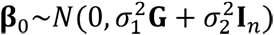 is the random intercept for individual *i*, accounting for population structure via the genetic relatedness matrix **G** and its associated variance 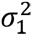, and the identity matrix **I**_***n***_ representing the individual-specific baseline level of saline self-administration for *n* individuals with variance 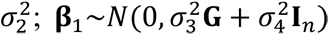 is the random slope of day for individual *i*, with associated variance 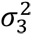 for genetic relatedness and variance 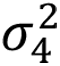 for individual day effect; 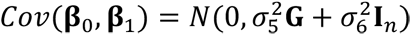 is the covariance between **β**_0_ and **β**_1_, with covariance 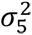 associated with the random intercepts and slopes attributable to **G**, and covariance 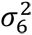 representing random intercepts and slopes not attributable to **G**; and *ε*_*ij*_∼*N*(0, *σ*^2^) is the residual error term.

Significance testing used Wald tests, which returned Z statistics, together with genome-wide significance thresholds of 5% obtained from permutation. After removing SNPs with minor allele frequency < 5% or missing genotype frequency > 10%, 340,097 remained [35]. Mouse genome build GRCm38/mm10 was employed [36].

#### 2.3.3 Open field

Comparison of saline IVSA endpoints and open field locomotor activity used data downloaded from the Mouse Phenome Database [37]. The open field measures were distance traveled in the first 10 min of a 55 min testing session for 62 inbred strains [38] and a 20 min testing session of 55 BXD recombinant inbred strains [39]. Distances were normalized as Z scores and the separate and combined datasets employed.

#### 2.3.4 GWASs of transcript, spliceform and RNA editing abundance

GWASs of transcript, spliceform and RNA editing abundance were performed as described using FaST-LMM to correct for population structure [10, 11, 13, 25]. Filtering of SNPs employed the same criteria as the behavioral traits. Behavioral cohort could not be used as a covariate for these analyses because RNA samples were barcoded and distributed over multiple sequencing lanes to minimize batch effects. Available covariates of sequencing batch and sex were used instead. *Cis* expression quantitative trait loci (eQTLs), splicing (percent spliced in, or ψ) QTLs (ψQTLs) and editing QTLs (ϕQTLs) were defined as residing within 2 Mb of the regulated gene. Behavioral or molecular QTLs were deemed coincident if located < 2 Mb apart [13].

#### 2.3.5 Transcriptome-wide association studies

`FUSION` and `FOCUS` software were used to perform transcriptome-wide association studies (TWASs) [26, 27]. Both methods incorporated sequencing batch and sex as covariates and returned Z statistics. Significance thresholds used *P* < 0.05, Bonferroni corrected for the number of genes tested. `FOCUS` is a Bayesian approach that also reports a posterior inclusion probability (pip), which was employed as the principal thresholding criterion with pip > 0.8.

#### 2.3.6 eCAVIAR

`eCAVIAR` was used to find single nucleotide polymorphisms (SNPs) that co-regulated *cis* eQTLs and behavioral loci. The analysis incorporated sequencing batch and sex as covariates. `eCAVIAR` is a Bayesian approach, and reports the highest colocalization posterior probability (CLPP) [24]. The CLPP represents the inclusion probability that a SNP co-regulates a gene and a trait. Markers within 200 SNPs of the *cis* eQTL were evaluated. The recommended support threshold for a co-regulating SNP is CLPP > 0.01, which displayed low false positive rates in multiple situations [24].

## 3 RESULTS

### 3.1 Saline intravenous self-administration

We evaluated 84 strains of the HMDP for IVSA of saline (477 mice) or cocaine (479 mice) over a 10 day testing period, as described previously [10]. Pressing one lever in the testing chamber caused delivery of the infusate (saline or cocaine, depending on experiment), while the other lever was inactive. Infusate delivery was accompanied by flashing of the house light and extinction of the aperture lights for 20 s. Four behavioral endpoints were evaluated: number of infusions, active lever presses, percent active lever presses and inactive lever presses.

The infusions represent the number of times saline was delivered as a result of active lever presses during the 2 h testing period. After delivery of the infusate, a 20 s time-out period was imposed during which active lever presses continued to be recorded, but no infusions delivered. Active lever presses are the sum of productive lever presses that deliver an infusion and non-productive presses during the time-out period. Infusions are thus a subset of active lever presses. Inactive lever presses represent the number of times the inactive lever, which has no programmed consequence, was pressed during the testing period. Percent active lever presses were calculated as the number of active lever presses divided by the sum of active and inactive lever presses.

Taken across the HMDP as a whole, there was a significant increase in normalized inactive lever presses as testing day increased (t[1,476] = 4.7, *P* = 3.4 × 10^-6^, Kenward-Roger df) and a significant decrease in percent active lever presses (t[1,476] = 6.1, *P* = 3.0 × 10^-9^) (Figure 1A). Neither infusions (t[1, 476] = 0.83, *P* = 0.41) nor active lever presses (t[1,476] = 0.22, *P* = 0.83) showed significant changes. Despite the overall trends, there were substantial interindividual differences between slopes and intercepts of the endpoints for individual mice. For example, mice showed dramatic interindividual differences in infusions, even though this endpoint showed no significant time dependence across all the mice (Figure 1B, Figures S1A,B).

**FIGURE 1.**
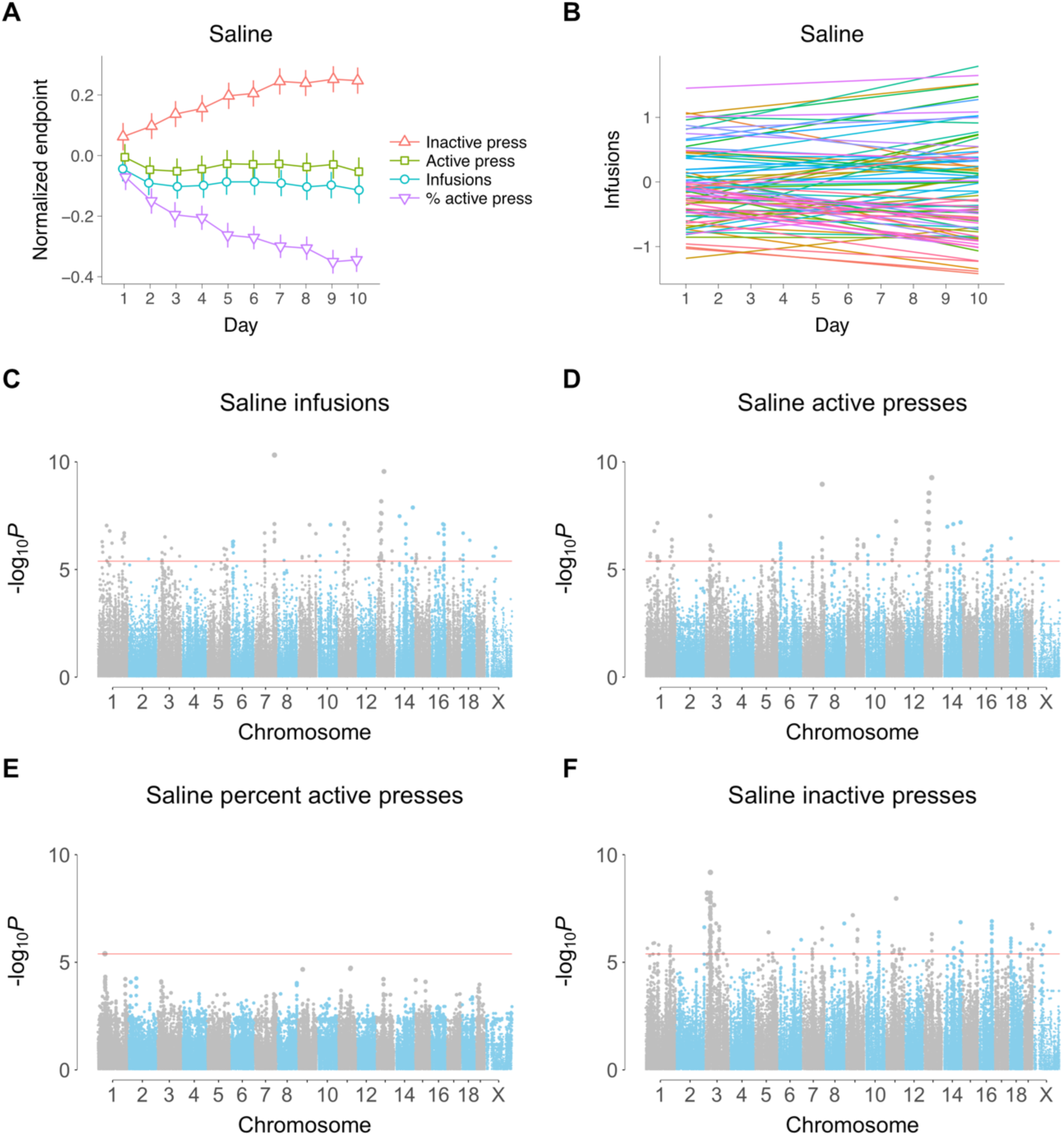
Intercepts, slopes and longitudinal genome scans for saline IVSA. (A) Time course for the four behavioral endpoints (means ± s.e.m.) (B) Intercept and slope of testing day for each of the 84 strains. Strains are plotted for clarity, since using individual mice gives a very dense plot. (C) Infusions GWAS. Each dot represents a SNP. Significance value indicated on ordinate (y axis) and genome coordinates for each SNP on abscissa (x axis). Red horizontal line, family-wise error rate = 5%. (D) Active lever presses. (E) Percent active lever presses, (F) Inactive lever presses.

Genome-wide association studies (GWASs) for saline IVSA using the four endpoints on each of the ten days produced no genome-wide significant loci [10, 11]. To improve fit and increase statistical power, we used `GMMAT` software to evaluate the longitudinal phenotypes via a linear mixed model (Figure 1B, Figures S1A,B) [28]. The day of assay was used as a continuous variable with both a fixed and random slope for individual mice together with a random intercept and random slope to correct for genetic relatedness, as described [10, 11]. We reasoned that this model would improve power, even for behavioral endpoints that showed no significant change with testing day, by correctly accounting for the substantial interindividual differences in time courses.

Due to the linkage disequilibrium structure of the HMDP, quantile-quantile (QQ) plots resulting from `GMMAT` showed inflation (Figures S1C-F). Percent active saline presses showed little inflation at highly significant *P* values, but still had a high genomic inflation factor because the bulk of the inflation was for more moderate *P* values tightly packed near the origin of the plot (Figure S1E).

A total of 145 significant loci, of which 85 were unique, were identified using the four behavioral endpoints of saline IVSA (Figure 1, Table S2). Of the 145 loci for saline IVSA, 56 were for infusions, 35 were for active lever presses, 1 for percent active lever presses and 53 for inactive lever presses. In contrast, 17 significant loci were obtained for cocaine using `GMMAT`, of which 15 were unique [11]. The greater number of loci for saline IVSA is consistent with its higher broad and narrow sense heritabilities.

The only locus significant for both cocaine and saline IVSA was for inactive lever presses on Chromosome 3 [11]. The peak SNP for saline was rs30114031 (37,799,968 bp, *P* = 6.6 × 10^-10^), which was more significant than the peak SNP for cocaine, rs30059671 (38,178,200 bp, *P* = 3.2 × 10^-7^). *Spry1* (sprouty RTK signaling antagonist 1) is centromeric to both SNPs, but closer to the saline SNP (157,696 bp distant) than the cocaine SNP (535,928 bp).

A triplet of loci for infusions, active lever presses, and inactive lever presses spanned from 66,206,986 bp to 68,567,223 bp (2,360,237 bp) on Chromosome 11 (Figure 2A). *Myh4* (myosin heavy chain 4) was located at the approximate center of this region (67,249,238 bp).

**FIGURE 2.**
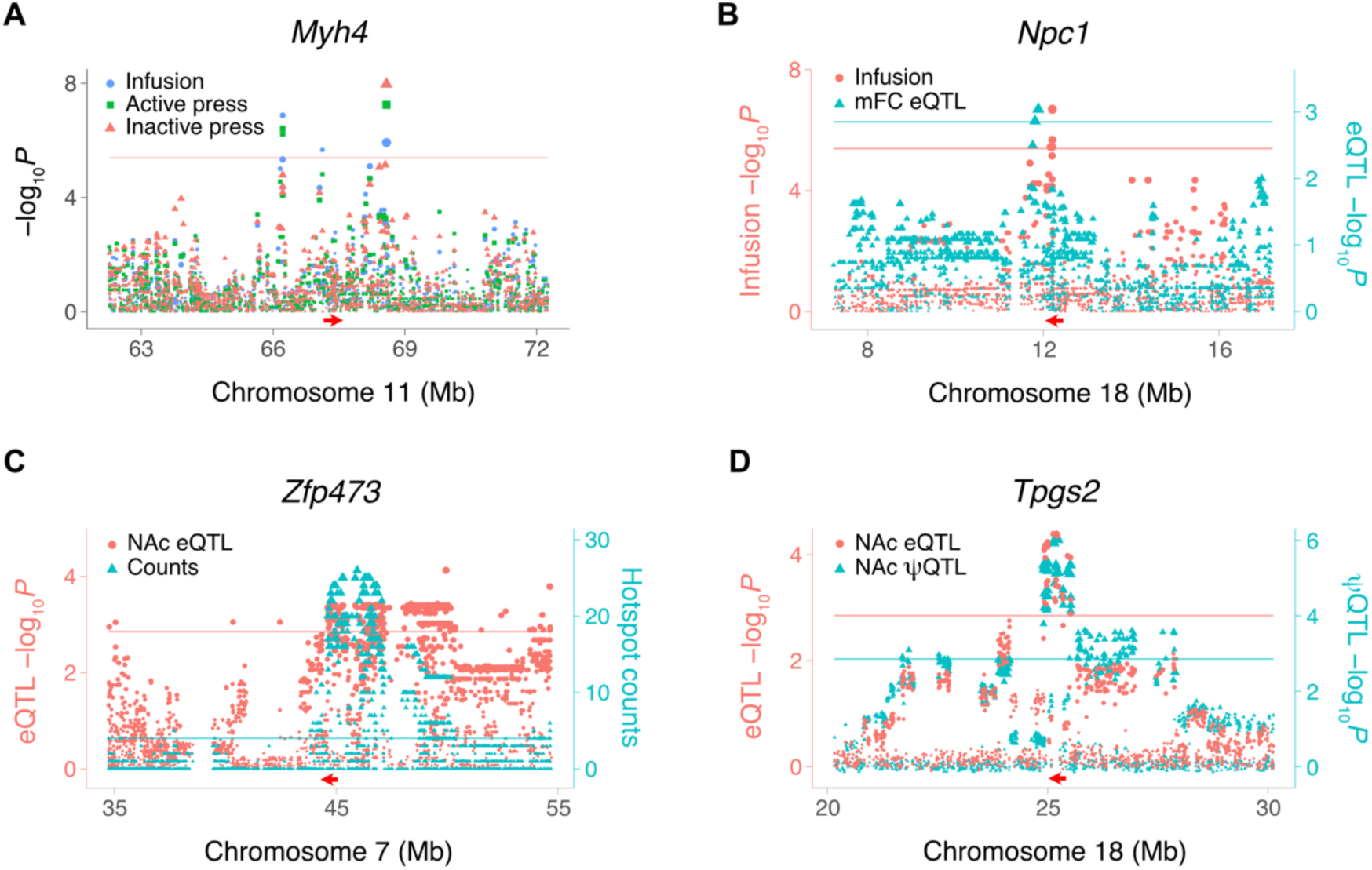
Loci for IVSA and gene expression in saline-exposed mice. (A) Myh4 is close to loci for three different saline IVSA endpoints. QTL peaks for infusions and active lever presses centromeric to Myh4 (left); infusions (center peak); infusions, active lever presses and inactive lever presses peaks telomeric to Myh4 (right). Horizontal red arrow, Myh4 location. Red horizontal line, family-wise error rate = 5%. Ordinate, infusion −log_10_P values. (B) Coincident loci for saline infusions and Npc1 cis eQTL in mFC. Left ordinate (red), infusion −log_10_P values. Right ordinate (blue), eQTL −log_10_P values. Red and blue horizontal lines, significance thresholds for infusions and cis eQTL, respectively. (C) Co-aligned NAc eQTL hotspot and Zfp473 cis eQTL. Red arrow, location of Zfp473. Left ordinate (red), eQTL −log_10_P values. Right ordinate (blue), number of genes regulated by each SNP. Red horizontal line, cis eQTL significance threshold. Blue horizontal line, hotspot count significance threshold, FDR < 0.05 (Poisson). (D) Coincident NAc cis eQTL for Tpgs2 and cis ψQTL for Tpgs2 exon 6. Peak marker rs31436205 for both QTLs. Left ordinate (red), eQTL −log_10_P values. Right ordinate (blue), ψQTL −log_10_P values. Red and blue horizontal lines, significance thresholds for cis eQTL and cis ψQTL, respectively.

### 3.2 RNA-Seq

To identify genetic pathways for saline IVSA, RNA-Seq was performed on NAc and mFC from 41 of the cocaine- and saline-exposed strains of the HMDP. NAc and mFC were chosen because of their role in operant self-administration of cocaine [3, 6]. A total of 73.0 ± 1.1 M paired-end reads were obtained per strain for saline, averaged over the two brain regions (NAc 72.2 ± 1.5 M; mFC 73.8 ± 1.5 M) [10]. Transcripts, spliceforms and RNA editing sites regulated by infusate, brain region and sex were previously discussed [11].

### 3.3 Expression QTLs

*Cis* and *trans* QTLs were mapped for transcript (expression QTLs, or eQTLs), splicing (sQTLs or ψQTLs) and RNA editing (edit QTLs or ϕQTLs) levels in the saline samples using `FaST-LMM` [25]. The number of *cis* eQTLs averaged over the two brain regions for saline was 4,878 ± 253 (NAc, 4,699; mFC, 5,057). The distance between the *cis* eQTLs and their corresponding gene was 0.62 Mb ± 0.009 Mb, averaged across brain regions (NAc, 0.63 ± 0.01 Mb; mFC, 0.62 ± 0.01 Mb), and is consistent with the linkage disequilibrium of the HMDP [12, 14, 40].

*Npc1* (Niemann-Pick disease, type C1 intracellular cholesterol transporter 1) showed a *cis* eQTL co-aligned with a QTL for infusions, suggesting a possible link between *Npc1* and this behavior (Figure 2B). Hotspots, in which a locus regulates many genes, were identified for transcript abundance [13, 15]. A total of 12 hotspots regulating ≥ 20 genes were present in NAc saline samples (FDR < 2.2 × 10^-16^), and 22 hotspots in mFC saline samples (FDR < 2.2 × 10^-16^). We sought candidate genes for hotspots by looking for co-aligned *cis* eQTLs. One NAc saline hotspot was coincident with a *cis* eQTL for the transcription factor *Zfp473* (zinc finger protein 473) (Figure 2C).

### 3.4 Splicing QTLs

A total of 1,418 ± 46 *cis* splicing QTLs (ψQTLs) were detected for saline exposed mice, averaged over the two brain regions (NAc, 1,385; mFC, 1,450). Transcript abundance can be affected by genetic variants that alter spliceform preference and hence mRNA stability [41, 42]. To evaluate the prevalence of this phenomenon, we examined whether there was a statistically significant enrichment in coincident *cis* eQTLs and *cis* ψQTLs. There were significant enrichments in both NAc and mFC. A total of 362 coincident *cis* eQTLs and ψQTLs were found in NAc from saline treated mice (odds ratio, OR, = 2.7, *P* < 2.2 × 10^-16^, Fisher’s exact test), and 398 in mFC (OR = 2.6, *P* < 2.2 × 10^-16^). Similar results were found for the cocaine exposed samples [11], indicating this enrichment is not infusate specific.

An example of a coincident *cis* eQTL and *cis* ψQTL for exon 6 of *Tpgs2* (tubulin polyglutamylase complex subunit 2) in NAc of saline exposed mice is shown in Figure 2D. The peak SNP, rs31436205, was the same for both the *cis* eQTL and *cis* ψQTL (Figure S2). The C allele was associated with higher percent spliced in of exon 6 of *Tpgs2* and with lower transcript abundance, consistent with inclusion of this exon destabilizing the mRNA.

### 3.5 Editing QTLs

Sequence changes caused by RNA editing can alter transcript stability and abundance as well as changing coding sequence [43]. A total of 262 ± 31 *cis*-acting loci were identified that affect RNA editing efficiency (ϕQTLs) in saline-exposed mice averaged over the two brain regions (NAc, 284; mFC, 240) [44].

Convincing quantitation of RNA editing event is more demanding than for transcript or spliceform abundance, since the editing causes single nucleotide changes that can decrease alignment rates. We defined the ascertainment rate of RNA editing events as the percent of samples for which alignment frequencies at an editing site exceeded the threshold of > 10% in each infusate and brain region. Editing events showed an ascertainment rate of 36.7 ± 0.4% across RNA-Seq samples from saline-exposed mice averaged over the two brain regions (NAc, 36.3 ± 0.3%; mFC, 37.0 ± 0.3%). The less than 100% detection rate implies decreased statistical power and indicates that the ϕQTLs should be treated with some caution. Increased read depth would help overcome this problem.

We looked for statistically significant enrichment in coincident *cis* eQTLs and ϕQTLs to evaluate how often genetically determined variations in RNA editing can change transcript levels. In NAc from saline treated mice, there were 57 co-aligned *cis* eQTLs and ϕQTLs (odds ratio, OR, = 2.1, *P* = 1.8 × 10^-6^, Fisher’s exact test) and 45 in mFC (OR = 1.8, *P* = 4.7 × 10^-4^, Fisher’s exact test), both significant enrichments. Co-aligned *cis* ϕQTLs and *cis* eQTLs should show significantly increased numbers of editing sites located in the mature mRNA compared to intronic or intergenic regions, assuming that *cis* ϕQTLs regulate editing events which in turn alter transcript stability and result in *cis* eQTLs. Indeed, we found significantly increased numbers of editing sites in 5’ untranslated, 3’ untranslated and coding regions compared to intronic and intergenic regions for the coincident *cis* ϕQTLs and eQTLs in both NAc (odds ratio = 2.0, *P* = 3.2 × 10^-3^, Fisher’s Exact Test) and mFC (odds ratio = 2.5, *P* = 2.0 × 10^-4^, Fisher’s Exact Test) of saline treated mice.

Statistical enrichment of coincident *cis* eQTLs and ϕQTLs, as well as a corresponding enrichment of editing sites in 5’ untranslated, 3’ untranslated and coding regions was also found in cocaine exposed mice [11], indicating that these phenomenon are not infusate specific.

### 3.6 Transcriptome-wide association studies

To pinpoint individual genes for saline IVSA, we employed transcriptome-wide association studies (TWASs). This approach acknowledges that most complex trait loci act through variation in expression regulatory sequences, and hence transcript levels, rather than amino acid sequence changes [45]. A gene for a trait is identified by TWAS when the genetically-predicted part of the gene’s expression is significantly correlated with the trait. TWAS enhances statistical power because the approach analyzes loci at the gene rather than marker level, leading to decreased multiple hypothesis correction. `FUSION` and `FOCUS` packages were employed to perform TWAS, with `FOCUS` providing additional fine mapping compared to `FUSION` [26, 27].

A total of 15 genes were uncovered using `FUSION` of saline exposed NAc, of which 9 were unique (Figure 3). Five genes were uncovered using saline exposed mFC, of which 4 were unique (Figure S3). There was no overlap between the NAc and mFC TWASs. A total of 5 unique genes were significant using `FOCUS` in NAc (posterior inclusion probability, pip > 0.8; active lever press: *Serpinb1a,* serpin family B member 1, pip = 0.97; *Pcnp*, PEST proteolytic signal containing nuclear protein, pip = 0.91; inactive lever press: *Cbx2*, chromobox 2, pip = 0.94) and in mFC (infusions: *5031434O11Rik*, pip = 0.95; active lever press: *5031434O11Rik*, pip = 0.90; inactive lever press: *Tmem241*, transmembrane protein 241, pip = 0.94). Two genes, *5031434O11Rik* and *Serpinb1a*, were significant using both `FUSION` and `FOCUS`.

**FIGURE 3.**
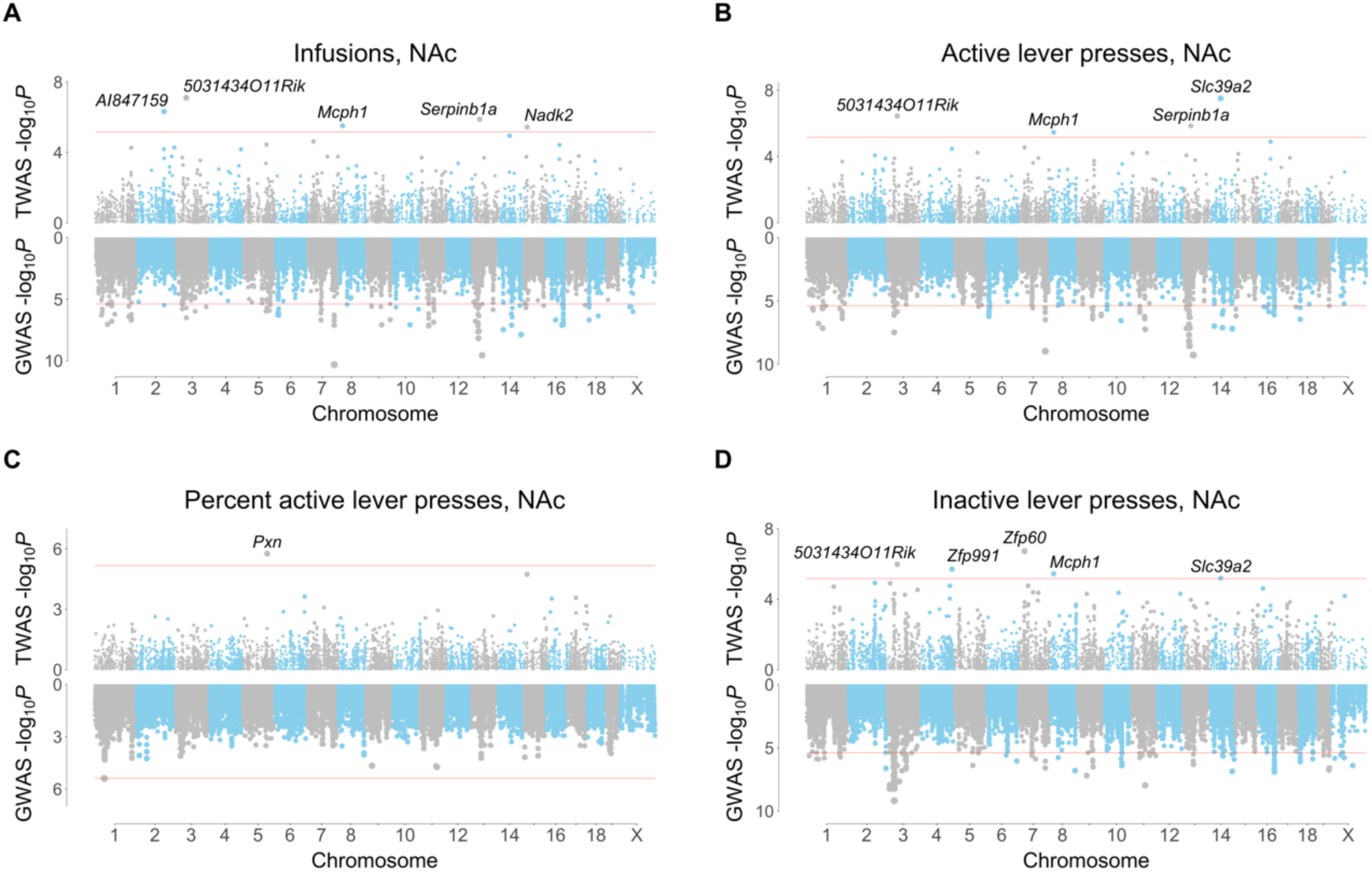
`FUSION` TWASs for saline IVSA using NAc RNA-Seq. (A) Infusions. (B) Active lever presses. (C) Percent active lever presses. (D) Inactive lever presses. TWASs on top, GWASs on bottom. Respective significance thresholds indicated by horizontal red lines.

`FUSION` identified *Zfp60* (zinc finger protein 60) as significant for inactive lever presses in NAc of saline exposed mice (Figure 3). Decreased *Zfp60* expression was associated with increased inactive lever presses. Another gene, *5031434O11Rik*, was significant for three out of four IVSA endpoints (infusions, active lever presses and inactive lever presses) using `FUSION` of saline exposed NAc (Figure 3). The gene was also significant for infusions (pip = 0.95, −log_10_*P* = 6.1) and active lever presses (pip = 0.90, −log_10_*P* = 5.7) using `FOCUS` of saline exposed mFC (Figure 4). Increased expression of *5031434O11Rik* was associated with increased lever presses.

**FIGURE 4.**
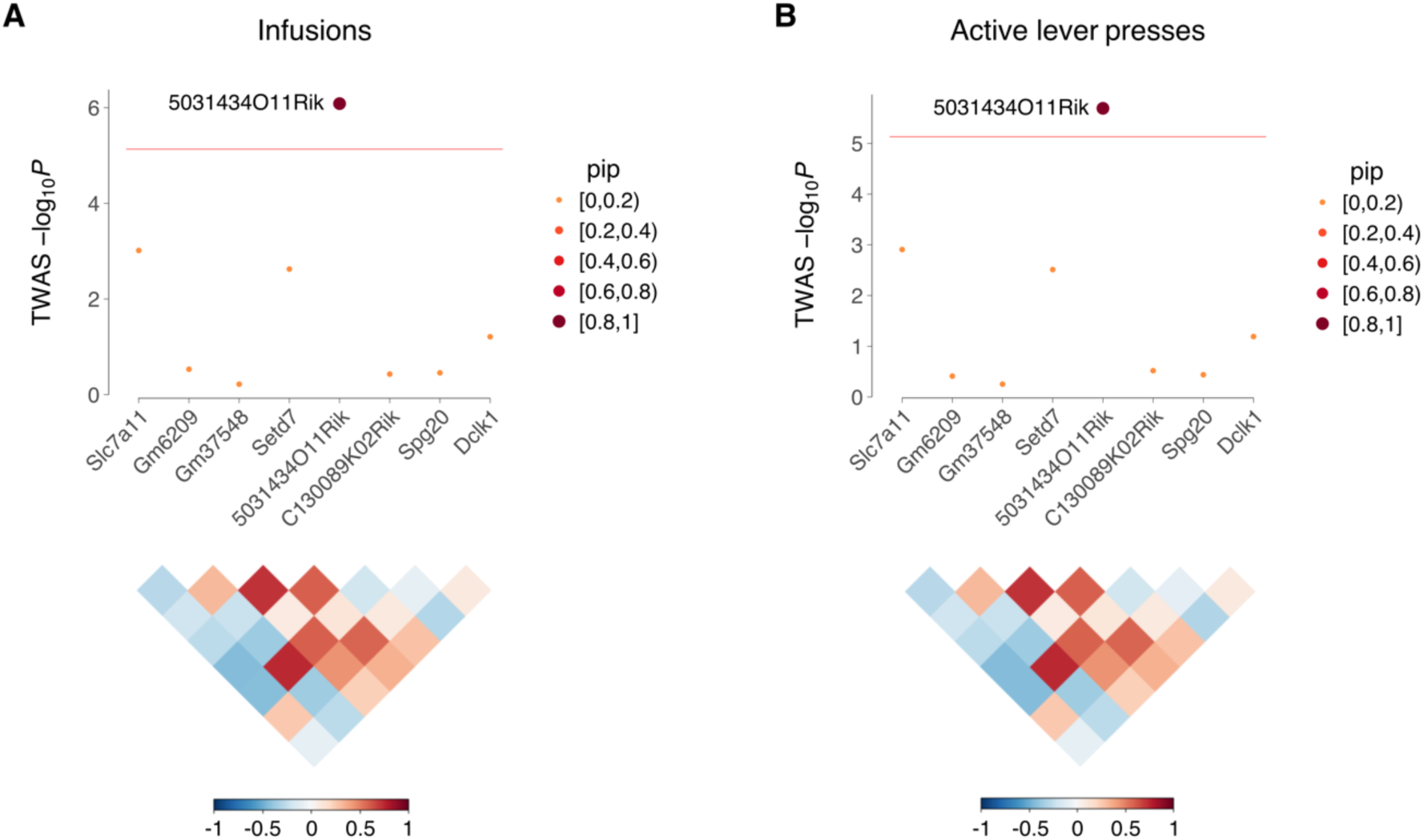
`FOCUS` of mFC in saline exposed mice. (A) `FOCUS` of infusions for region on Chromosome 3 containing 5031434O11Rik. The ordinate shows −log_10_P calculated from `FOCUS` TWAS Z scores. Horizontal red line indicates transcriptome-wide significance threshold. Point sizes indicate pip. Correlation coefficients (R) of predicted expression values between genes, together with corresponding color scale, shown below. (B) `FOCUS` of 5031434O11Rik region for active lever presses. For both endpoints, pip of 5031434O11Rik ≥ 0.9. For all other genes, pip < 0.2.

The peak SNP for infusions near to *5031434O11Rik* on Chromosome 3 was rs49204785 at 52,563,394 bp, located 999,818 bp telomeric to the gene (Figure 5A) [33]. The C allele of rs49204785 was associated with significantly higher saline infusions compared to the T allele (effect size = 0.40 ± 0.08, *P* = 3.1 × 10^-7^) (Figure 5B). Consistent with the effect of *5031434O11Rik* being saline specific, there was no significant allelic difference of rs49204785 for cocaine.

**FIGURE 5.**
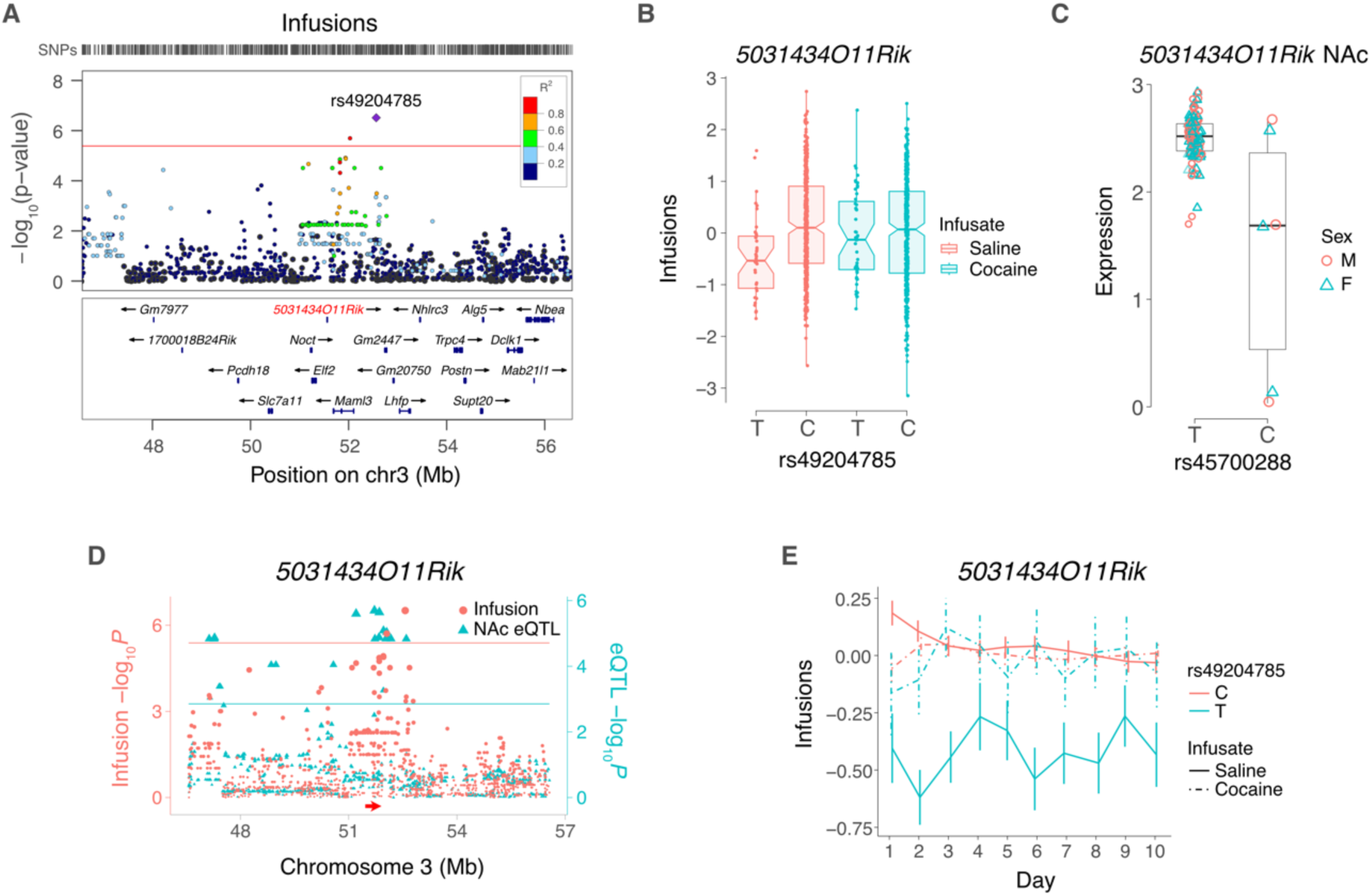
5031434O11Rik and saline infusions. (A) LocusZoom plot for saline infusions, showing locus harboring 5031434O11Rik (red). Peak SNP is rs49204785. R^2^ values indicate linkage disequilibrium between peak and adjacent SNPs. (B) The peak behavior SNP shows significant allelic effect on normalized infusion endpoint for saline but not cocaine. (C) Normalized expression of 5031434O11Rik in saline NAc depends on allele of peak SNP, rs45700288, for 5031434O11Rik eQTL. (D) Coincident loci for saline infusions (left ordinate, red) and 5031434O11Rik cis eQTL in saline NAc (right ordinate, blue). Red and blue horizontal lines, significance thresholds for infusions and cis eQTL, respectively. (E) Infusion time course for peak behavior SNP of 5031434O11Rik locus, rs49204785, with saline or cocaine infusate. Means ± s.e.m.

The peak SNP for the *5031434O11Rik cis* eQTL was rs45700288 (51,711,018 bp), which was in significant linkage disequilibrium with rs49204785 (D′ = 0.83, *R*^2^ = 0.63, df = 475, *P* < 2.2 × 10^-16^) (Figures 5C,D). The T allele of rs49204785 showed significantly decreased saline infusions (interaction between SNP and infusate: Z = 4.3, *P* = 2.0× 10^-5^) (Figures 5E). In contrast, the differences between the effects of the C allele on saline infusions and either allele on cocaine infusions were less significant (Z < 2.5, *P* > 0.01).

Although rs45700288, the peak SNP significant *cis* eQTL for *5031434O11Rik*, exceeded the thresholds of minor allele frequency > 5% and missing genotype frequency < 10%, only 3 strains had the minor allele in the RNA-Seq data (SJL/J, NZW/LacJ and BXD98/RwwJ) (Figures 5C,D and Figure S4). However, each datapoint represents an average of pooled samples from 3 individuals (male or female) increasing confidence in the conclusions.

### 3.7 eCAVIAR

We used `eCAVIAR` to further evaluate the contribution of *5031434O11Rik* to saline IVSA. `eCAVIAR` accounts for the uncertainty introduced by linkage disequilibrium while evaluating the posterior probability that both GWAS and eQTLs are caused by the same SNP [24]. Support for the same causal variant is given by a colocalization posterior probability (CLPP) > 0.01.

Consistent with a causal role for saline IVSA suggested by the TWASs, `eCAVIAR` identified rs48952099 (52,050,163 bp on Chromosome 3) as the peak suprathreshold SNP for the *5031434O11Rik cis* eQTL and behavioral endpoints (infusions, NAc CLPP = 0.016; mFC CLPP = 0.018; active lever presses, NAc CLPP = 0.015; mFC CLPP = 0.016). This SNP was roughly half-way between the peak eQTL SNP (rs45700288; 51,711,018 bp) and the peak SNP for saline infusions (rs49204785; 52,563,394 bp).

The *5031434O11Rik* gene encodes a long non-coding (lnc) RNA gene and overlaps with, and is a potential antisense transcript to, the 5′ end of a neighboring gene, *Setd7* (SET domain containing 7, histone lysine methyltransferase) [46]. To test whether *5031434O11Rik* acts by destabilizing the *Setd7* transcript, we evaluated the relationship between *5031434O11Rik* and *Setd7* transcript levels in the combined saline NAc and mFC samples using a linear mixed model with tissue and strain as random effects. There was no significant relationship between *5031434O11Rik* and *Setd7* mRNA abundances (t[1,108] = 1.5, *P* = 0.14). Similarly, there was no significant correlation between the transcript levels in an RNA-Seq atlas of mouse tissues using tissue as a random effect (t[1,32] = 1.9, *P* = 0.07) [47].

### 3.8 Saline IVSA and open field

Percent active lever presses were used in an attempt to normalize IVSA based on locomotor activity. Of the 145 loci for saline IVSA, only one was significant for percent active lever presses. In addition, only one gene was significant for percent lever presses using `FUSION`, `FOCUS` and `eCAVIAR`. The paucity of loci for percent lever presses suggested that the loci for the other saline IVSA endpoints may be detecting genetic effects for phenotypes related to spontaneous locomotor activity. To test this conjecture, we compared the saline IVSA endpoints with distance traveled in the open field for inbred and BXD recombinant inbred strains [37–39]. There were no significant correlations using the combined inbred and BXD datasets or the inbred dataset alone. However, there were some weak positive correlations between the saline IVSA endpoints and the BXD dataset alone (infusions, *R* = 0.32, df = 40, *P* = 0.04; active lever presses, *R* = 0.32, df = 40, *P* = 0.04).

## 4. DISCUSSION

We previously used longitudinal linear mixed models to identify 17 behavioral loci for cocaine IVSA in the HMDP [10, 11]. In this study, we used the same approach to analyze a parallel group of control mice that had undergone concurrent saline IVSA. We found 145 loci for the saline IVSA, only one of which overlapped with cocaine. The smaller heritability and decreased number of detected QTLs for cocaine compared to saline indicates that cocaine IVSA has a larger environmental contribution and is due to loci with smaller average effect sizes. Full genetic understanding of cocaine use disorder will likely continue as an enduring problem.

Infusions and active lever presses showed no significant change with testing day. In contrast, inactive lever presses showed a highly significant increase, suggesting reinforcement. The reinforcing stimulus is puzzling, since pressing the inactive lever offers no obvious contingent reward such as an infusion or flashing of the house lights. The increasing number of inactive lever presses also contrasts with spontaneous locomotor activity which typically decreases over testing day [48]. Unsurprisingly, percent active lever presses showed a highly significant decrease with day of testing, since this normalized endpoint is calculated by dividing active lever presses by total lever presses (active plus inactive lever presses).

We included the X chromosome in the GWASs for saline IVSA and found 5 unique loci (2 for infusions and 3 for inactive lever presses). GWASs of the X chromosome are complicated by a number of factors, including copy number differences in males and females, X chromosome inactivation in females, and altered recombinations and Hardy-Weinberg equilibria compared to the autosomes [49, 50]. Because of these complications, geneticists have been reluctant to report X chromosome data in human GWASs, leading to pleas to include the chromosome [49, 50]. The situation is simpler in the HMPD, since the mouse resource employs genetically stable strains. In addition, our analyses employed sex as a covariate, helping to alleviate problems due to pleiotropy and the other confounds. Nevertheless, it is wise to treat our X chromosome loci with care.

Four potential genes emerging from the saline IVSA analyses, *Myh4*, *Npc1*, *Zfp60* and *5031434O11Rik*, were implicated in locomotor activity by previous studies [46, 51–54]. One region on Chromosome 11 harbored three loci that regulated saline infusions, active lever presses and inactive lever presses. In the middle of this region was *Myh4*, a myosin heavy chain gene. In a previous study, an intronic SNP in *Myh4* was positively selected in mouse lines bred for voluntary wheel running [51]. This SNP causes the mini-muscle phenotype which, depending on body mass, results in significantly reduced quadriceps and gastrocnemius muscle masses, as well as larger soleus muscles [55, 56]. Mini-muscle individuals had faster but decreased duration of wheel running. The genetic regulation of saline IVSA could thus be due to non-neural as well as neural effects.

*Npc1* possessed a *cis* eQTL that was coincident with a QTL for saline infusions, suggesting a possible role for the gene in this behavior. *Npc1* was also significant in a human GWAS study for walking pace (*P* = 3 × 10^-11^) [52, 53].

The long non-coding (lnc) RNA gene, *5031434O11Rik*, was implicated in saline infusions, active lever presses and inactive lever presses using `FUSION` TWAS, and in infusions and active lever presses using `FOCUS` TWAS and `eCAVIAR`. The *5031434O11Rik* gene is expressed in the olfactory region, cortex, hippocampus and cerebellum [57]. A recent study found that *5031434O11Rik* was the most strongly differentially expressed gene in the striatum of four mouse lines selectively bred for high voluntary wheel running compared to non-selected controls [46]. Increased expression of *5031434O11Rik* was associated with decreased activity, opposite to our TWAS results. The different genetic backgrounds in the wheel running and present studies may explain the contrasting relationship.

Compared to protein coding genes, our understanding of lncRNA genes is poor, although a role in brain development is emerging [58, 59]. Many lncRNA genes regulate chromatin, supporting the idea that *5031434O11Rik* may be an antisense transcript that destabilizes the overlapping mRNA of its neighboring gene, *Setd7* [46]. However, we found no significant relationship between *5031434O11Rik* and *Setd7* transcript levels, suggesting that if *5031434O11Rik* acts by regulating chromatin it uses a mechanism other than destabilization of *Setd7* transcripts.

Another gene emerging from the TWASs was *Zfp60*. A knockout of this zinc finger gene showed significant hyperactivity in multiple endpoints of both the open-field and light-dark tests (*P* = 8.8 × 10^-9^ and 1.2 × 10^-7^, respectively) [54]. *Zfp60* is expressed in olfactory bulb, hippocampus, cortical subplate and hypothalamus based on *in situ* hybridization, and regions related to operant conditioning, such as midbrain, striatum, and pallidum, based on single cell RNA-Seq [57, 60]. Apart from the mouse studies of *5031434O11Rik* and *Zfp60*, genes identified for saline IVSA in the TWASs showed no significant associations for locomotor-related phenotypes in a human GWAS catalog [52].

Only one locus was significant for percent active lever presses out of the 145 saline loci and only one gene was significant for this endpoint using `FUSION`, `FOCUS` and `eCAVIAR`. The `FUSION` significant gene for percent lever presses, *Pxn* (paxillin), encodes a protein that is part of the post-synaptic adhesome and may play a role in learning and memory, a behavior linked to drug use disorder [61, 62]. Since percent lever presses are a measure of saline IVSA normalized to total lever presses, the lack of loci for this endpoint are consistent with the other endpoints being surrogate measures of locomotion. However, there was only a weakly significant relationship between two saline IVSA endpoints (infusions and active lever presses) and distance traveled in the open field. Any connections between the saline IVSA and locomotor phenotypes could be clarified by analysis of larger datasets. The relative absence of significant loci for percent active lever presses also suggests that the loci for the other three endpoints are unlikely to be due to noise, and that the *P* value threshold is appropriate for genome-wide association.

Significant enrichment of ψQTLs and ϕQTLs with their co-aligned *cis* eQTLs was found in both saline exposed NAc and mFC. In addition, coincident *cis* ϕQTLs and eQTLs showed significant enrichment of editing sites in untranslated regions and coding regions compared to intronic or intergenic regions. Splicing or RNA editing can thus alter transcript abundance by changing mRNA stability. Similar results were found in the cocaine IVSA study, suggesting that this phenomenon is general.

The saline IVSA study was designed as a control for cocaine and to dissect the genetic similarities and differences between these behaviors. Indeed our results indicate distinct genetic foundations for the two traits. However, the physiological relevance of saline IVSA is unknown. In outbred rats that successfully acquired cocaine self-administration, operant responding reinforced only by a visual stimulus was correlated with subsequent lever pressing for cocaine [63]. However, our results suggest that reinforcement supplied by the saline infusion and flashing of the house light is regulated by partially different genetic pathways than that provided by cocaine. Notably, pressing the inactive lever does not flash the house light. The apparently reinforcing properties of inactive lever presses together with the large number of loci for this behavior, some in common with infusions and active lever presses, suggests that the light flash is not the only motivating factor contributing to lever pressing in saline IVSA.

Although some genes for saline IVSA also regulated locomotor activity, the connection between these traits was only weakly supported by open field data. Nevertheless, our work establishes that a control infusate for self-administration has minimal genetic overlap with a drug prone to misuse, and raises interesting questions about what exactly is being measured by saline IVSA.

## Author Contributions

Conceived and designed the study: EE, JDJ, DJS. Acquired the data: AHK, JRB. Analyzed the data: AHK, JRB, NL, CG-F, TCS, MC, XX, EE, JDJ, DJS. Wrote the paper: AHK, JRB, JDJ, DJS.

## Acknowledgements

We thank the UCLA Semel Institute Neurosciences Genomics Core for sequencing. This work used computational and storage services associated with the Hoffman2 Shared Cluster provided by the UCLA Institute for Digital Research and Education Research Technology Group.

## Data Availability Statement

The sequencing data generated in this study can be downloaded from the NCBI BioProject database (https://www.ncbi.nlm.nih.gov/bioproject/) under accession number PRJNA755328. Data and code are also available from figshare (https://figshare.com/; https://doi.org/10.6084/m9.figshare.27958836).

## Ethics Statement

Animal experiments were approved by the Binghamton University Institutional Animal Care and Use Committee. All procedures were conducted in accordance with guides on the humane care and use of laboratory animals issued by the National Institute of Health and the American Association for Accreditation of Laboratory Animal Care.

## Additional Information

Supplementary information is available.

