## Supplementary Information for "Differing genetics of saline and cocaine self-administration in the hybrid mouse diversity panel"

[7] Current address: Department of Pharmaceutical Sciences, Binghamton University, Binghamton, NY 13902

[8] Current address: Department of Human Genetics, University of Chicago, Chicago, IL 60637

[9] Current address: Sanford Burnham Prebys, La Jolla, CA 92037

**Table S1. Intravenous self-administration sample sizes**

| Strain | Saline | | | Cocaine | | | RNA-Seq |
|---|---|---|---|---|---|---|---|
| | M | F | Total | M | F | Total | |
| 129S1/SvImJ | 3 | 3 | 6 | 3 | 3 | 6 | None |
| 129X1/SvJ | 3 | 3 | 6 | 3 | 3 | 6 | Same sex pooled |
| A/J | 3 | 3 | 6 | 3 | 3 | 6 | Individual |
| AKR/J | 3 | 3 | 6 | 3 | 3 | 6 | Individual |
| BALB/cByJ | 3 | 3 | 6 | 3 | 3 | 6 | Same sex pooled |
| BALB/cJ | 4 | 3 | 7 | 4 | 3 | 7 | Same sex pooled |
| BPL/1J | 3 | 3 | 6 | 3 | 3 | 6 | Same sex pooled |
| BTBR T<+> Itpr3<tf>/J | 3 | 3 | 6 | 3 | 4 | 7 | None |
| C3H/HeJ | 3 | 3 | 6 | 3 | 3 | 6 | Same sex pooled |
| C3HeB/FeJ | 3 | 3 | 6 | 3 | 3 | 6 | Same sex pooled |
| C57BL/10J | 3 | 4 | 7 | 3 | 3 | 6 | Same sex pooled |
| C57BL/6J | 3 | 3 | 6 | 4 | 3 | 7 | Same sex pooled |
| C57BLKS/J | 3 | 3 | 6 | 3 | 3 | 6 | Same sex pooled |
| C57BR/cdJ | 3 | 3 | 6 | 3 | 3 | 6 | Same sex pooled |
| C57L/J | 5 | 3 | 8 | 3 | 3 | 6 | None |
| C58/J | 3 | 3 | 6 | 3 | 3 | 6 | Same sex pooled |
| CBA/J | 4 | 3 | 7 | 3 | 3 | 6 | Same sex pooled |
| DBA/1J | 3 | 3 | 6 | 3 | 3 | 6 | Same sex pooled |
| DBA/2J | 3 | 3 | 6 | 3 | 3 | 6 | Same sex pooled |
| FVB/NJ | 3 | 3 | 6 | 3 | 3 | 6 | Same sex pooled |
| I/LnJ | 2 | 2 | 4 | 1 | 2 | 3 | None |
| KK/HlJ | 3 | 3 | 6 | 3 | 3 | 6 | Same sex pooled |
| LP/J | 3 | 3 | 6 | 3 | 3 | 6 | Individual |
| MA/MyJ | 3 | 3 | 6 | 3 | 3 | 6 | Same sex pooled |
| MRL/MpJ | 3 | 3 | 6 | 4 | 3 | 7 | Same sex pooled |
| NOD/ShiLtJ | 4 | 3 | 7 | 3 | 3 | 6 | Individual |
| NZB/BlNJ | 2 | 3 | 5 | 4 | 3 | 7 | Same sex pooled |
| NZO/HlLtJ | 4 | 3 | 7 | 3 | 2 | 5 | Same sex pooled |
| NZW/LacJ | 3 | 3 | 6 | 3 | 3 | 6 | Same sex pooled |
| PL/J | 3 | 3 | 6 | 3 | 3 | 6 | Same sex pooled |
| SJL/J | 3 | 3 | 6 | 3 | 4 | 7 | Same sex pooled |
| SM/J | 3 | 3 | 6 | 3 | 4 | 7 | Same sex pooled |
| BXD1/TyJ | 4 | 3 | 7 | 3 | 3 | 6 | None |
| BXD2/TyJ | 2 | 3 | 5 | 3 | 3 | 6 | None |
| BXD6/TyJ | 2 | 1 | 3 | 2 | 2 | 4 | None |
| BXD9/TyJ | 3 | 3 | 6 | 3 | 3 | 6 | None |
| BXD11/TyJ | 2 | 3 | 5 | 3 | 3 | 6 | None |
| BXD13/TyJ | 2 | 3 | 5 | 3 | 3 | 6 | None |
| BXD14/TyJ | 3 | 2 | 5 | 2 | 1 | 3 | None |
| BXD15/TyJ | 3 | 3 | 6 | 2 | 3 | 5 | None |
| BXD16/TyJ | 3 | 3 | 6 | 3 | 2 | 5 | None |
| BXD18/TyJ | 3 | 2 | 5 | 2 | 2 | 4 | None |
| BXD19/TyJ | 3 | 3 | 6 | 3 | 3 | 6 | None |
| BXD21/TyJ | 3 | 1 | 4 | 3 | 3 | 6 | None |
| BXD27/TyJ | 3 | 2 | 5 | 3 | 2 | 5 | None |
| BXD28/TyJ | 3 | 3 | 6 | 2 | 3 | 5 | None |
| BXD29/TyJ | 2 | 2 | 4 | 2 | 3 | 5 | None |
| BXD31/TyJ | 3 | 3 | 6 | 4 | 3 | 7 | Same sex pooled |
| BXD32/TyJ | 3 | 3 | 6 | 3 | 3 | 6 | Same sex pooled |
| BXD33/TyJ | 2 | 3 | 5 | 1 | 3 | 4 | None |
| BXD34/TyJ | 3 | 4 | 7 | 3 | 4 | 7 | None |
| BXD38/TyJ | 3 | 3 | 6 | 3 | 3 | 6 | Same sex pooled |
| BXD39/TyJ | 3 | 3 | 6 | 3 | 3 | 6 | None |
| BXD40/TyJ | 3 | 3 | 6 | 3 | 2 | 5 | Same sex pooled |
| BXD42/TyJ | 3 | 3 | 6 | 3 | 3 | 6 | Same sex pooled |
| BXD43/RwwJ | 3 | 3 | 6 | 3 | 3 | 6 | None |
| BXD48a/RwwJ | 3 | 3 | 6 | 3 | 3 | 6 | Same sex pooled |
| BXD49/RwwJ | 1 | 2 | 3 | 3 | 2 | 5 | None |
| BXD50/RwwJ | 3 | 3 | 6 | 3 | 3 | 6 | None |
| BXD51/RwwJ | 3 | 2 | 5 | 2 | 2 | 4 | None |
| BXD55/RwwJ | 3 | 3 | 6 | 2 | 3 | 5 | None |
| BXD56/RwwJ | 2 | 3 | 5 | 2 | 3 | 5 | None |
| BXD60/RwwJ | 4 | 3 | 7 | 3 | 3 | 6 | None |
| BXD61/RwwJ | 3 | 3 | 6 | 3 | 3 | 6 | Same sex pooled |
| BXD62/RwwJ | 3 | 3 | 6 | 3 | 3 | 6 | Same sex pooled |
| BXD63/RwwJ | 3 | 3 | 6 | 2 | 3 | 5 | None |
| BXD65/RwwJ | 3 | 3 | 6 | 3 | 3 | 6 | Same sex pooled |
| BXD68/RwwJ | 3 | 3 | 6 | 3 | 3 | 6 | None |
| BXD69/RwwJ | 1 | 3 | 4 | 2 | 4 | 6 | None |
| BXD70/RwwJ | 3 | 3 | 6 | 3 | 3 | 6 | None |
| BXD71/RwwJ | 2 | 3 | 5 | 3 | 3 | 6 | None |
| BXD73a/RwwJ | 3 | 3 | 6 | 3 | 3 | 6 | Same sex pooled |
| BXD75/RwwJ | 3 | 2 | 5 | 2 | 3 | 5 | None |
| BXD77/RwwJ | 3 | 3 | 6 | 3 | 3 | 6 | Same sex pooled |
| BXD83/RwwJ | 1 | 3 | 4 | 3 | 3 | 6 | None |
| BXD84/RwwJ | 3 | 3 | 6 | 3 | 3 | 6 | Same sex pooled |
| BXD85/RwwJ | 2 | 3 | 5 | 3 | 3 | 6 | None |
| BXD86/RwwJ | 3 | 3 | 6 | 3 | 3 | 6 | None |
| BXD89/RwwJ | 3 | 1 | 4 | 3 | 1 | 4 | None |
| BXD90/RwwJ | 3 | 2 | 5 | 3 | 3 | 6 | None |
| BXD98/RwwJ | 3 | 3 | 6 | 3 | 3 | 6 | Same sex pooled |
| BXD99/RwwJ | 1 | 3 | 4 | 2 | 2 | 4 | None |
| BXD100/RwwJ | 3 | 3 | 6 | 3 | 3 | 6 | None |
| BXD102/RwwJ | 3 | 0 | 3 | 2 | 0 | 2 | None |
| **Mean ± s.e.m.** | **2.9 ± 0.07** | **2.8 ± 0.06** | **5.7 ± 0.10** | **2.9 ± 0.06** | **2.9 ± 0.07** | **5.7 ± 0.10** | |

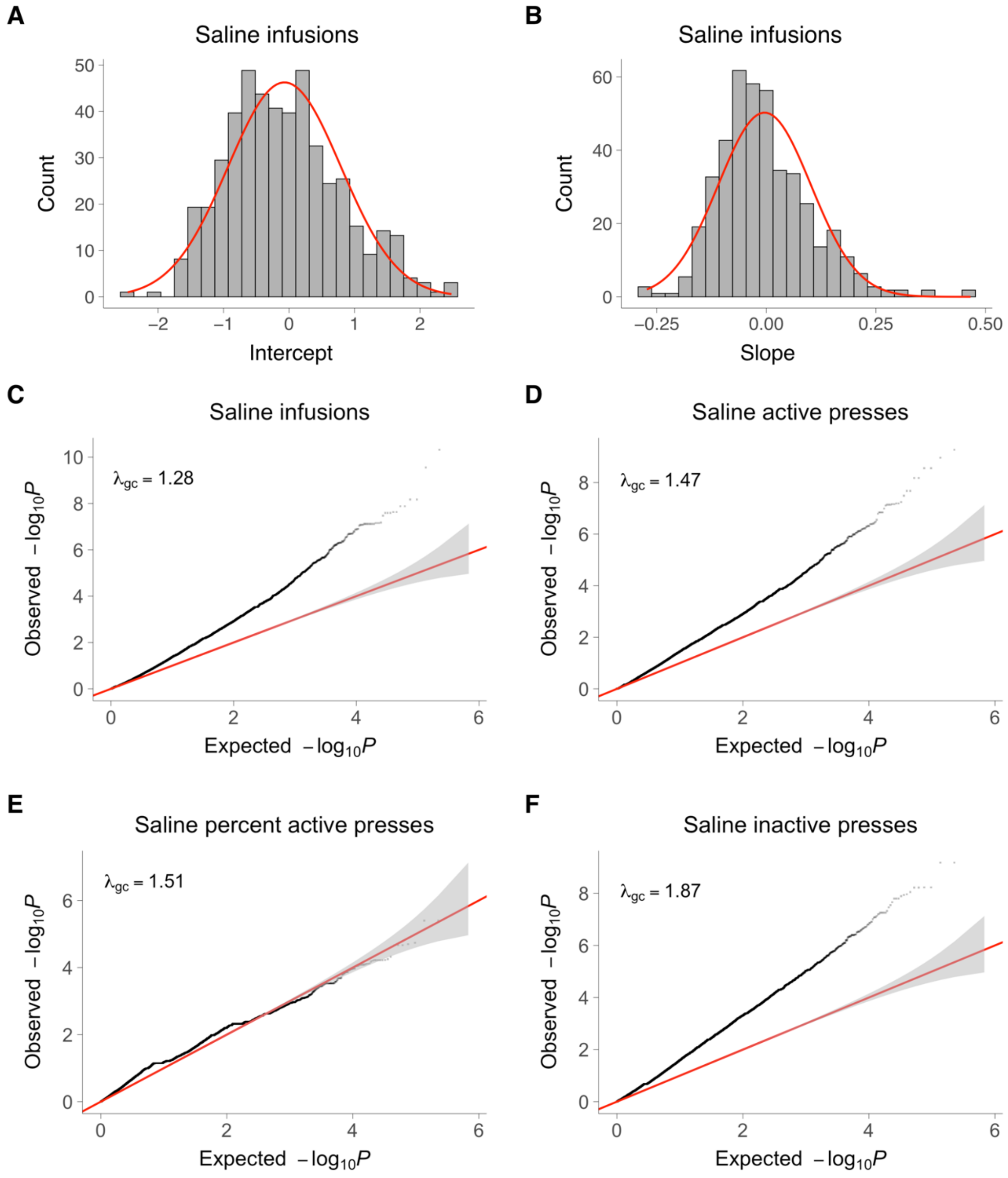


FIGURE S1 Intercepts, slopes and quantile-quantile (QQ) plots for longitudinal saline genome scans. (A) Infusion intercepts for individual mice (gray histogram bars). Normal distribution (red). (B) Infusion slopes for individual mice. (C) QQ plots for infusions. QQ plot for normal distribution (red). Grey area, 95% confidence interval. $\lambda_{gc}$, genomic control (inflation) factor. (D) Active lever presses. (E) Percent active lever presses. (F) Inactive lever presses.

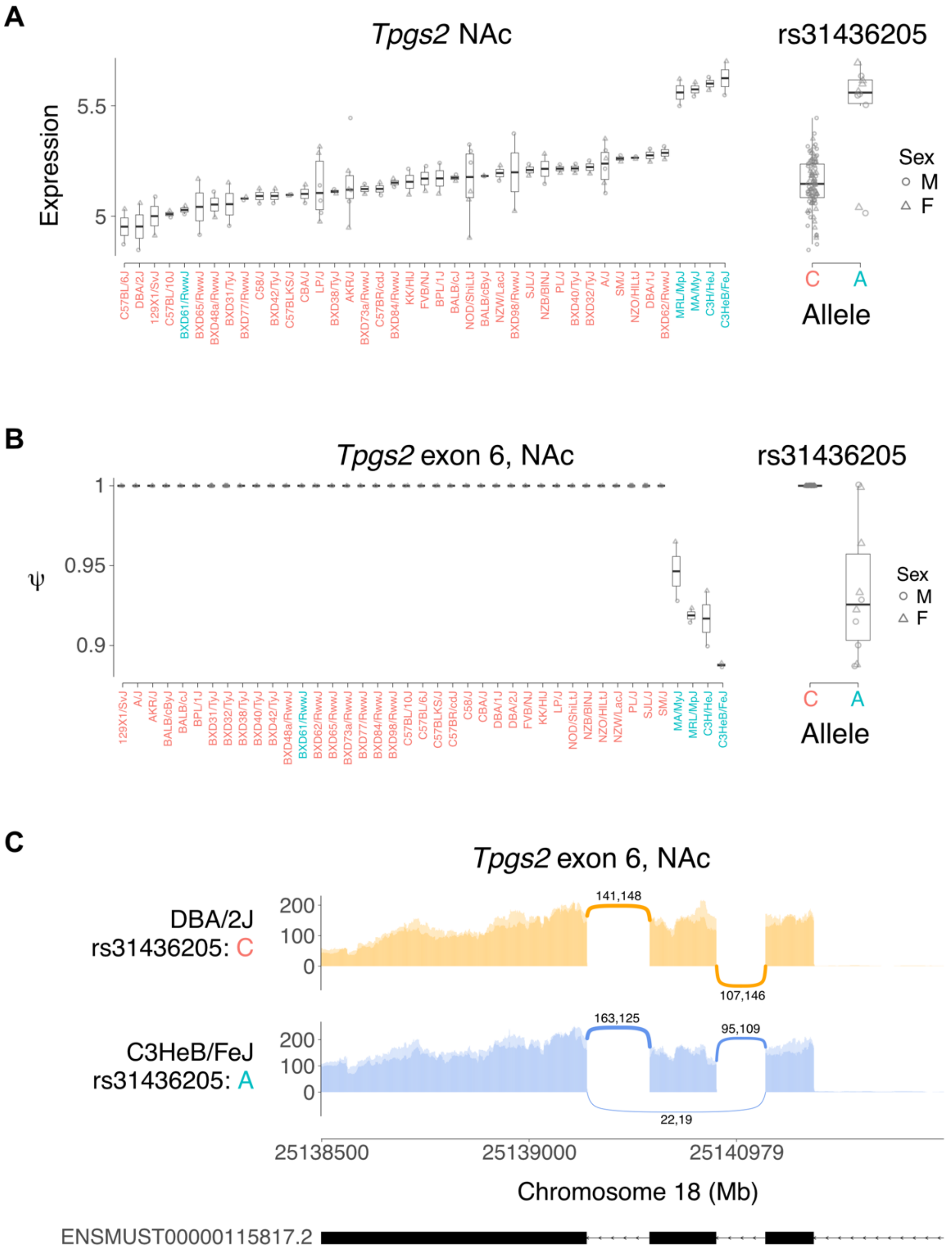


FIGURE S2 Genetic variation in alternate splicing of exon 6 *Tpgs2* in saline treated mice. (A) Strain survey of *Tpgs2* expression in saline NAc. Peak eQTL for *Tpgs2* is rs31436205. Strains with C allele tend to have low expression (red), while strains with A allele have high expression (blue). (B) Percent spliced in (psi or ψ) for exon 6 of *Tpgs2*. Strains with C allele have close to 100% ψ, while strains with A allele have lower ψ. (C) Sashimi plot [1] showing high ψ for *Tpgs2* exon 6 in DBA2/J, a strain with C allele of rs31436205, and low ψ for C3HeB/FeJ, a strain with A allele.

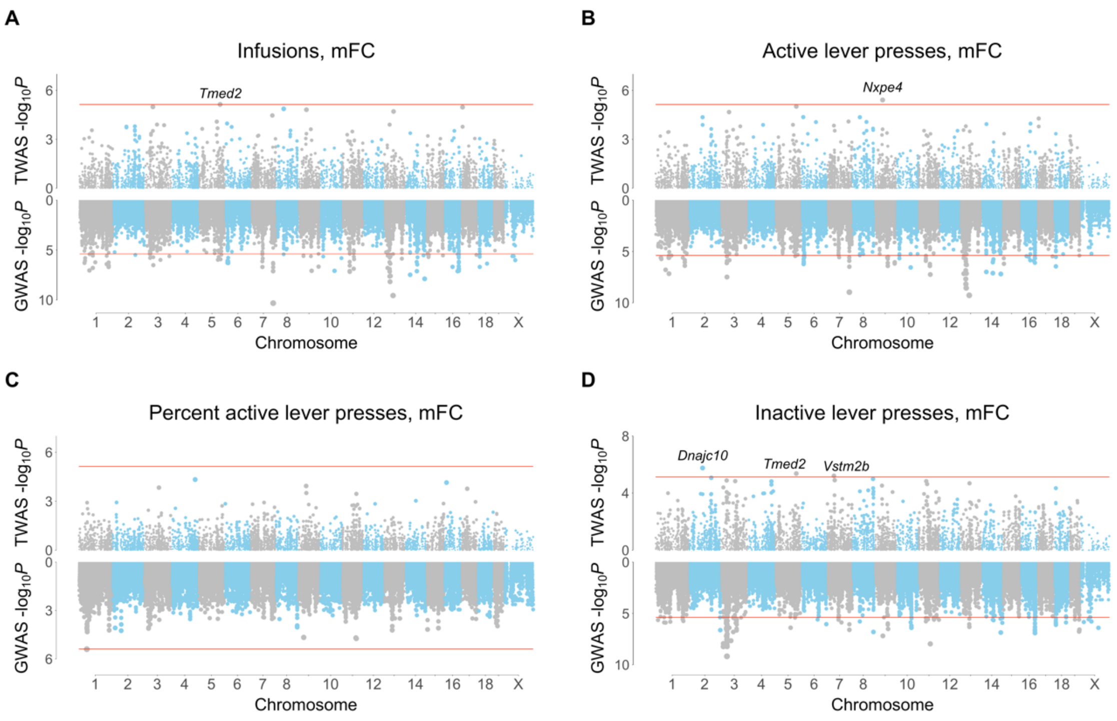


FIGURE S3 FUSION TWASs for saline IVSA using mFC RNA-Seq. (A) Infusions. (B) Active lever presses. (C) Percent active lever presses. (D) Inactive lever presses. TWASs on top, GWASs on bottom, with respective thresholds indicated by horizontal red lines.

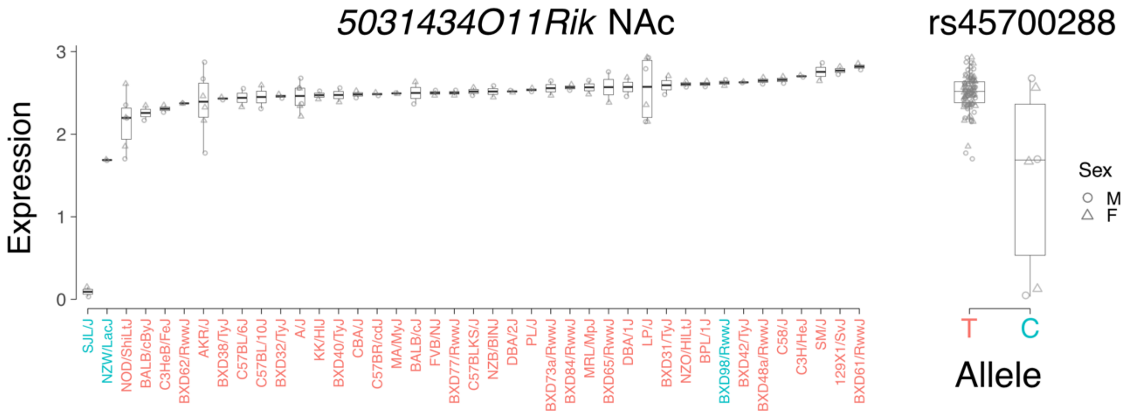


FIGURE S4 Expression of *5031434O11Rik* in saline treated mice. Strain survey of *5031434O11Rik* expression in NAc (left). Peak eQTL for *Tpgs2* is rs45700288 (right). Strains with T allele have high expression (red), while strains with the (minor) C allele have low expression (blue). Each datapoint for the C allele of rs45700288 (right) represents an average of pooled samples from 3 individuals (male or female).
